# Cerebellar function remains resilient under increased task demands in healthy adults up to 80 years but it is task-specific and independent of cerebellar structure

**DOI:** 10.64898/2026.04.02.716060

**Authors:** Anouck Matthijs, Anda de Witte, Dante Mantini, Jean-Jacques Orban de Xivry

## Abstract

Healthy aging is associated with progressive structural brain decline, yet the loss of functional abilities varies across individuals, which has been linked to reserve mechanisms. Within the framework of complex systems theory, reserve is thought to manifest as resilience when the system is challenged by stressors, such as increases in task difficulty. The cerebellum has been proposed as a potential source of motor reserve, but empirical evidence linking cerebellar structure, function, and resilience remains limited. We conducted a cross-sectional study including 50 young, 80 older, and 30 older-old adults to examine resilience to increasing task demands across cerebellar-specific and general outcomes. Participants completed three motor tasks (pure elbow motion, motor timing, postural stability) and two cognitive tasks (mental rotation, spatial working memory). Structural MRI was acquired to quantify cerebellar grey matter volume within functionally defined regions.

Cerebellar-specific motor measures (anticipatory muscle activation and timing variability) were preserved across age groups and remained resilient under increased task demands, including in adults over 80 years of age. In contrast, general sensorimotor performance (postural sway) declined with age and showed reduced resilience. Within the cognitive domain, both cerebellar-specific and general measures showed comparable age-related declines and reduced resilience. Resilience measures were not correlated across tasks, indicating that resilience is task- and domain-specific. Furthermore, cerebellar grey matter volume did not predict resilience in motor or cognitive outcomes.

These findings support the cerebellar motor reserve hypothesis, suggesting that cerebellar-dependent motor processes remain resilient despite age-related structural decline. However, resilience appears to be function-specific rather than a generalized individual trait. Overall, the results highlight dissociations between brain structure, function, and resilience, underscoring the selective contribution of the cerebellum to motor preservation in healthy aging.

## Introduction

Healthy aging is accompanied by gradual changes in brain and body systems that affect functional capacity in later life, including declines in motor, cognitive, and sensorimotor performance (Hunter et al., 2016; Seidler et al., 2010; Yang et al., 2023). As populations worldwide continue to age, preserving functional capacity has become a major societal and clinical priority, because preserved autonomy in older age not only benefits the individual’s quality of life but also reduces the pressure on healthcare systems.

Importantly, aging trajectories vary considerably across individuals: whereas some older adults experience pronounced functional decline, others maintain high levels of performance well into advanced age (Hunter et al., 2016; Prince et al., 2024; Yaffe et al., 2009; Zimmermann et al., 2025). These inter-individual differences often become most apparent when individuals are required to perform tasks under increased demands or in the presence of stressors (Kok et al., 2020). For example, two individuals may walk comfortably at the same speed under low-demand conditions, yet adopt markedly different walking speeds when confronted with a more challenging environment, such as an uneven or unstable surface. In gerontological research, exposure to stressors encompasses both long-term challenges, such as declines in physical health, social isolation, and bereavement (Smith & Hayslip, 2012), and short-term stressors, including acute illness, daily hassles, or sudden disruptions to routine, all of which can adversely affect functioning and well-being (Gagnon & Wagner, 2016; Leger et al., 2018; Whitehead & Torossian, 2021). Understanding the mechanisms underlying these inter-individual differences in responses to stressors is therefore crucial for developing strategies to promote healthy aging and sustain independent living in an aging society.

An individual’s response to a stressor provides an indication of their resilience, reflecting the system’s intrinsic capacity to withstand and adapt to challenges (Chhetri et al., 2021). Intrinsic capacity broadly refers to the composite of an individual’s physical and mental capacities and is characterized by five interacting domains: locomotion, cognition, vitality, psychological function, and sensory function (Beard et al., 2016; Chhetri et al., 2021). Together, these domains capture overall functional capacity. Intrinsic capacity may further be viewed as a higher-level manifestation of underlying reserve processes (Chhetri et al., 2021). Reserve is commonly used to explain how individuals maintain functional abilities despite age-related biological decline (such as brain atrophy), reflecting latent neural or physiological resources that help buffer performance. Accordingly, individuals with greater reserve and intrinsic capacity are expected to exhibit more resilient responses when confronted with functional challenges.

Measuring individuals’ resilience could substantially improve our understanding of age-related differences and inform strategies to promote healthy aging. However, resilience is a latent construct that can only be inferred in the presence of a stressor (Links et al., 2018), posing a methodological challenge. Many real-life stressors relevant to aging (e.g., hip fracture, loss of a loved one) occur unpredictably, leaving researchers little control over their timing and magnitude and making resilience difficult to estimate directly. Consequently, resilience must typically be assessed indirectly using carefully controlled and ethically acceptable experimental paradigms.

The complex systems approach provides a framework for studying resilience in experimental settings by conceptualizing organisms as complex systems whose functional stability can be evaluated by introducing controlled stressors and how the system respond to them (Gijzel et al., 2019; Koivunen et al., 2024). Within this framework, resilience can be quantified using stimulus-response-based indicators (Varadhan et al., 2008). A stimulus refers to a controlled stressor applied to the system, whereas the response reflects the system’s measurable reaction to that stressor. For example, systematically increasing the difficulty of a motor task can serve as a standardized stressor, and the resulting change in performance (e.g., accuracy, stability, or timing precision) reflects the system’s response to it. When the applied stressor is relevant to the targeted system under investigation, the resulting stimulus–response relationship, such as how strongly performance deteriorates or is maintained under increasing task demands, may serve as a proxy for real-life resilience of the individual’s system (Kok et al., 2020; Varadhan et al., 2008). Stimulus–response paradigms therefore allow systems to be perturbed safely and in a controlled manner, while enabling resilience to be quantified through the characterization of response dynamics, including resistance to the stressor, recovery processes, and fatigability.

This framework also enables the assessment of resilience both between individuals and across different subsystems within the same individual. In this context, subsystems may refer to neural networks or functional control processes supporting specific behaviors, such as motor coordination, motor timing, or postural stability. Because functional outcomes emerge from interactions among these subsystems and resilience across subsystems is assumed to contribute to the functioning of the individual as a whole (Scheffer et al., 2018), resilience observed in one task may therefore relate to resilience in other tasks. However, age-related declines in resilience are unlikely to occur uniformly across the body (Scheffer et al., 2018). Instead, resilience may be differentially preserved across subsystems, such that some may remain relatively robust and partially compensate for declines in more vulnerable systems (Scheffer et al., 2018). Identifying such relatively preserved subsystems may therefore help reveal sources of reserve capacity that support overall individual resilience.

A potentially promising source of reserve capacity for motor-related processes is the cerebellum. The cerebellum plays a central role in motor control through the formation of internal models that enable predictive regulation of movement, making it essential for accurate movement planning and execution (Manto et al., 2012; Shadmehr & Krakauer, 2008; Wolpert & Ghahramani, 2000). Structurally, cerebellar volume is known to decline with age (Raz et al., 2005). Consequently, several studies have proposed that age-related cerebellar atrophy contributes directly to declines in motor control (Bernard & Seidler, 2014; Hogan, 2004). However, converging evidence suggests that cerebellar function may be relatively preserved even in advanced age. Experimental studies examining motor tasks that rely on cerebellar-specific mechanisms, including grip–load force coupling (Gilles & Wing, 2003), inter-joint coordination (Lee et al., 2007; Matthijs et al., 2025), sensory attenuation (Wolpe et al., 2016), motor adaptation (Vandevoorde & Orban de Xivry, 2019), and motor timing (Duchek et al., 1994), have often reported no age-related impairments. Furthermore, a recent study of de Witte et al. (2026b) demonstrated that adults up to 80 years of age maintained cerebellar-specific motor performance despite measurable reductions in cerebellar volume, whereas more general motor outcomes did exhibit age-related decline (de Witte et al., 2026b). Collectively, these findings support the notion that the cerebellum may constitute a particularly important source of motor reserve, demonstrating preserved functional outcomes despite age-related structural degeneration (Elbaz et al., 2013; Fleischman et al., 2015).

Despite accumulating evidence suggesting that the cerebellum may serve as a source of motor reserve, key questions remain unresolved. First, it is unclear whether resilience of cerebellar function represents a general property of the individual or whether it is specific to particular tasks or functional domains. Although the cerebellum contributes to both motor and cognitive processes (Arleo et al., 2023; Clark et al., 2021; Stoodley, 2012), most existing evidence focuses on motor function, leaving it unclear whether age-related resilience of cerebellar function extends beyond motor functions to cognitive domains and whether it generalizes across different task outcomes. Second, previous studies have rarely examined cerebellar function under systematically manipulated task demands. Because resilience can only be revealed when a system is sufficiently challenged, experimental paradigms that probe performance across levels of difficulty are essential for quantifying resilience. Furthermore, it has been demonstrated that cerebellar structure does not predict cerebellar-specific task outcomes (de Witte et al., 2026b). However, it remains unclear whether individual differences in resilience under such challenging task conditions are related to cerebellar structure. Addressing these gaps is critical to advancing our understanding of cerebellar reserve and its role in adaptive functioning during aging.

The present study was designed to address these gaps by investigating whether resilience of cerebellar function is maintained under increased task demands and whether it is restricted to the motor domain or generalizes across tasks and functional domains. To this end, we assessed resilience across three motor task, two targeting cerebellar-specific processes and one reflecting general sensorimotor function, and two cognitive tasks, one targeting cerebellar-specific cognitive process and one targeting general cognitive function. Consistent with the complex system approach, in each task we incorporated a task-specific stressor to systematically increase task demands. This design enabled the characterization of the stimulus-response relationship, providing an index of individual resilience for each task. To complement these behavioral measures, all participants underwent structural MRI of the whole brain, allowing us to examine whether greater cerebellar gray matter volume could predict enhanced resilience-related functional outcomes across the different tasks. To ensure that age-related effects were captured across the adult lifespan, our participant sample included participants over 80 years of age.

Based on the cerebellar motor reserve hypothesis (Elbaz et al., 2013; Fleischman et al., 2015), we hypothesize that cerebellar-specific motor outcomes would be maintained across aging, even under increased task demands, and that this resilience would be partly independent of age-related structural decline of the cerebellum. In contrast, we expect cerebellar-specific cognitive outcomes, as well as general motor and cognitive measures, to exhibit less resilience to stressors in older adults. Finally, we hypothesize that resilience across tasks would be positively associated, particularly between the two motor tasks targeting cerebellar-specific processes, suggesting that resilience of cerebellar function may reflect a broader individual characteristic rather than a purely task-specific phenomenon. By integrating behavioral indices of resilience with structural neuroimaging across motor and cognitive domains, this study aims to clarify the role of the cerebellum in task outcomes during healthy aging.

## Methods

### 2.1. Participants

A total of 161 individuals participated in the study. Participants were distributed across three age groups: 50 young adults (YA; 20–35 years, mean ± SD = 23.3 ± 2.0), 80 older adults (OA; 55–70 years, mean ± SD = 63.9 ± 4.6), and 31 older-old adults (OOA; 80+ years, mean ± SD = 82.4 ± 1.6). Data from one participant in the oldest group were excluded prior to analysis due to non-compliance with task instructions. Recruitment was conducted as part of a larger cross-sectional project examining age-related changes in cerebellar motor function. The overall sample size was determined based on previous work assessing structural cerebellar differences across these age ranges (Walhovd et al., 2011), which reported large group effects (Cohen’s d > 0.65; YA vs. OA: 0.98; OA vs. OOA: 0.66). Such effect magnitudes were considered adequate to capture the corresponding functional differences. Power calculations indicated that 50 participants in each of the young and older groups and 30 participants in the older-old group would yield 80% power to detect such effect sizes. To support prospective longitudinal follow-up of the older group, its size was increased to 80 participants, allowing for the detection of small effects (r = 0.3) with 80% power while accounting for an expected attrition rate of 20% (Bell et al., 2013).

All participants were screened for right-handedness using the Edinburgh Handedness Inventory (Oldfield, 1971) and for cognitive functioning via the Montreal Cognitive Assessment (MoCA), with a minimum score of 23 required for inclusion (Carson et al., 2018). Health screening confirmed that all participants were free from neurological or psychiatric conditions, substance dependence, stroke, and smoking. Additionally, participants with contraindications to MRI were not eligible for this study, with the exception of participants in the oldest age group. All participants provided written informed consent prior to participation in the experiment. The study protocol was approved by the KU Leuven/UZ Leuven ethics committee (project S66650).

### 2.2. Paradigm

All participants completed two experimental sessions of approximately 2.5 hours each. Across sessions, participants performed a range of tasks assessing cerebellar motor function, general motor performance, proprioception, cognition, and physical fitness. The present study is part of this broader cross-sectional study and therefore focuses only on three motor tasks: (1) the pure elbow motion task, (2) the motor timing task, and (3) the postural stability task; and on two cognitive tasks: (1) the mental rotation task and (2) the spatial working memory task. Across both sessions, task order was fixed to minimize fatigue effects. Cognitively demanding tasks were scheduled at the start of a session or immediately following a short break to ensure optimal engagement. Specifically, the spatial working memory task was performed first in the first session, while the mental rotation task occurred approximately one hour into the second session, preceded by a five-minute break. The pure elbow motion task was always performed at the beginning of the second session, as it required the longest preparation time (∼30 min). The postural stability task was completed halfway through the first session (after ∼1 hour), also following a short break. Finally, the motor timing task was performed at the end of either the first or second session, depending on age group, approximately two hours after the start of the session.

Each of these tasks was designed with a task-specific stressor to enable the assessment of resilience in motor and cognitive control mechanisms under increased task demands, following the complex systems theory of resilience (Scheffer et al., 2018). Within both the motor and cognitive domains, outcome measures were categorized as either cerebellar-specific or general sensorimotor or cognitive measures. This approach allowed us to examine whether resilience represents a cerebellum-specific property or a more general cross-domain capacity. Below, the task protocols are described, together with the classification of each outcome measure as cerebellar-specific or general. **Table 1** provides an overview of all outcome measures and the number of participants included in the analysis for each task.

**Table 1:** Overview of each task, including the task domain, outcome measure, number of included participant per age group, and reasons for participant exclusion (if applicable).

| Task name | Task domain | Outcome measure | Number of included participants |  |  | Reasons for exclusion |
| --- | --- | --- | --- | --- | --- | --- |
| Pure-elbow motion | Motor | Anticipatory muscle timing<br>(Cerebellar-specific measure) | YA = 32<br>(64 %) | OA = 50<br>(63 %) | OOA = 24<br>(80 %) | Problems with data saving (n = 6 YA, 14 OA, 2 OOA);<br>EMG onset not found (n = 12 YA, 16 OA, 4 OOA) |
| Motor timing | Motor | Clock variance<br>(Cerebellar-specific measure) | YA = 48<br>(96 %) | OA = 80<br>(100 %) | OOA = 29<br>(97 %) | Inattentive during experiment (n = 2 YA, 1 OOA) |
| Postural balance | Motor | Postural sway<br>(General measure) | YA = 50<br>(100 %) | OA = 79<br>(99 %) | OOA = 30<br>(100 %) | Problem of the lower limbs (n = 1 OA) |
| Mental rotation | Cognitive | Reaction time<br>(Cerebellar-specific measure) | YA = 50<br>(100 %) | OA = 80<br>(100 %) | OOA = 30<br>(100 %) |  |
| Spatial working memory | Cognitive | % False answers<br>(General measure) | YA = 50<br>(100 %) | OA = 80<br>(100 %) | OOA = 30<br>(100 %) |  |
| MRI |  | Cerebellar grey matter volume | YA = 49<br>(98 %) | OA = 78<br>(98 %) |  | Data lost (n = 1 YA);<br>Not eligible for MRI (n = 2 OA) |

### 2.3. Motor tasks

#### **Pure elbow motion task** (cerebellar specific outcome measure: anticipatory muscle activation)

The pure elbow motion task was performed while participants were seated in the KINARM exoskeleton (KINARM, BKIN Technologies Ltd., Kingston, ON, Canada), with surface EMG electrodes (Delsys Trigno Avanti Sensors, Boston, MA, USA) recording muscle activity from the right (dominant) arm. Participants performed 30° pure elbow flexion and extension movements. The task has been described in detail in a previous study (Matthijs et al., 2025). That study showed no directional differences in anticipatory agonist timing between flexion and extension movements. Therefore, the current study was limited to a single movement direction. Extension was selected because valid EMG data was available for the largest number of participants for this movement direction. During the elbow movement, vision of the hands and arms was occluded, and hand position was represented by a cursor displayed in an augmented reality setup. To isolate feedforward control strategies, visual feedback of hand position was removed during movement execution. Elbow extension movements always started with the shoulder at 60° (external angle) and the elbow at either 105° or 75° (external angles) (**Figure 1A**). Participants were instructed to perform a pure rotation of the forearm around the elbow while keeping the shoulder stable and to stop the movement as close as possible to the target. During such multi-joint movements, rotation at the elbow joint generates additional forces at the shoulder joint because of the limb’s inertia. To execute pure elbow rotations accurately, the nervous system must predict these interaction torques and anticipate shoulder muscle activity to counteract them and stabilize the shoulder joint. This anticipatory activation becomes increasingly important when interaction torques increase, such as at higher movement speeds.

**Figure 1:**
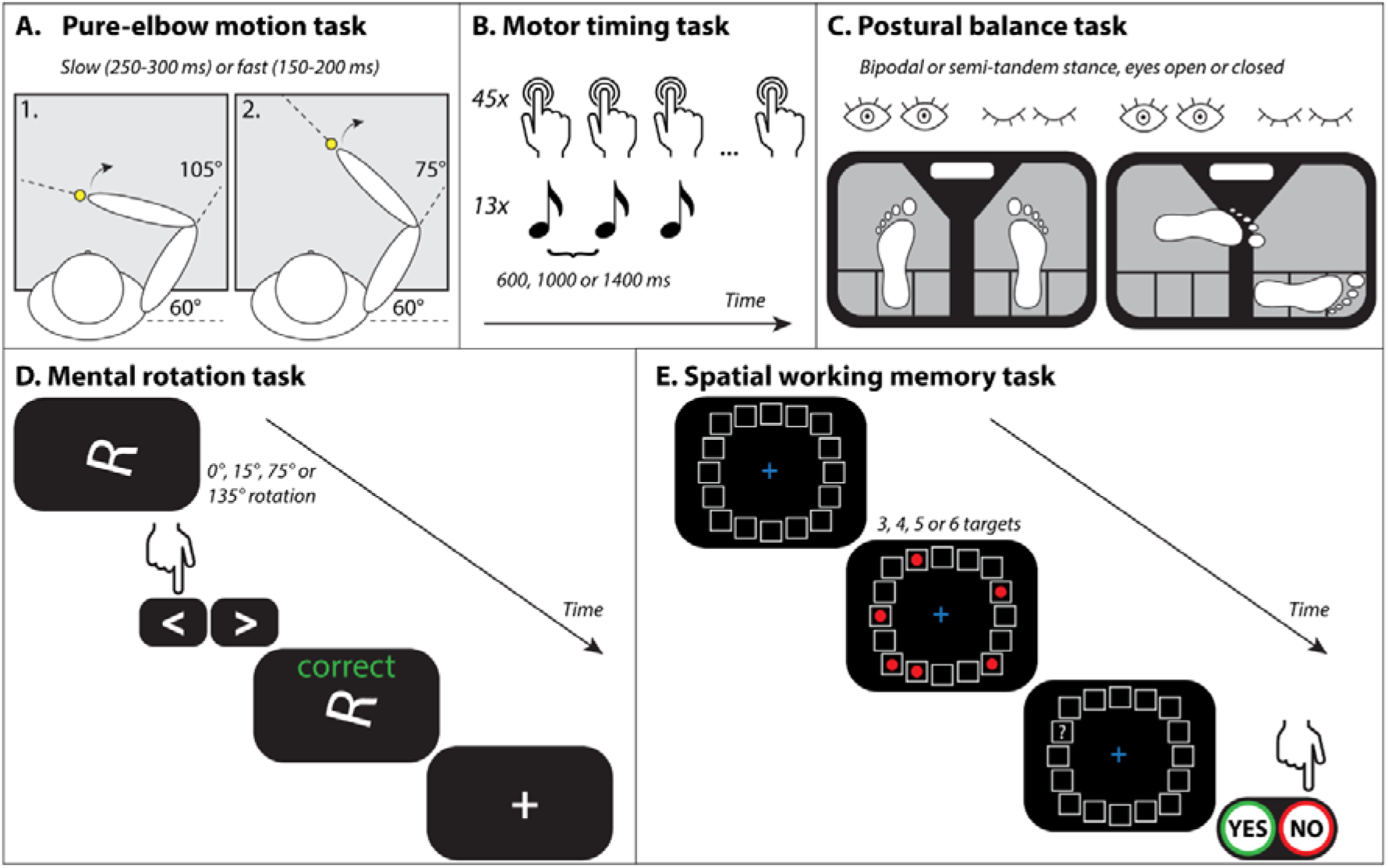
Overview of all different tasks and the included stressor in each task. **A**) Pure elbow motion task: 2 start positions of the upper arm from which participants performed 30° extension elbow movements in two speed conditions (slow: 250-300 ms; fast: 150-200 ms). The target is positioned in such a way that it can be reached while keeping the shoulder at 60° (external angle). **B**) Motor timing task: Participants start by tapping the space bar of a computer in sync with the beeps (interval of 600, 1000, or 1400 ms between beeps). The beeps cease after 13 taps, but the participant keeps tapping the same rhythm as accurately as possible for an additional 31 taps without any feedback. **C**) Postural balance task: visualization of the two possible foot positions on the balance plate (Btracks), with and without visual feedback. Participants are positioned with their hand on their hips and fixate on a cross 1-2 m in front of them. The position is held for 35 seconds, and a beep indicates the start and end of one trial. **D**) Mental rotation task: overview of the sequence of one trial. A letter stimulus in white was presented on a black background and remained visible until the response. Participants responded by pressing the corresponding arrow on the computer as fast as possible (left arrow: standard form, right arrow: mirrored). Feedback after each trial was provided, followed by a fixation cross. **E**) Spatial working memory: overview of the sequence of one trial. After a blank grid is displayed for 1 second, three to six targets appear at random positions within the circular grid for 2 seconds. A question mark then appears at one location in the grid, and participants have 5.5 seconds to indicate “yes” or “no” by pointing the KINARM robot to report whether a red target was previously present at that location.

Therefore, to investigate whether anticipatory activation could be maintained under increased interaction torques, participants performed the task under two controlled speed conditions in separate blocks: (1) slow (250–300 ms) and (2) fast (150–200 ms). Movement duration was calculated for each trial as the time from leaving the start position to reaching the target. Increasing movement speed was considered the task-specific stressor, as it increases the intersegmental interaction torque at the shoulder and thereby places greater demands on predictive motor coordination between the shoulder and elbow joints. After each trial, participants received color-coded feedback (green = correct speed, blue = too slow, orange = too fast) to help maintain the desired movement speed. The order of the slow and fast speed conditions was counterbalanced across participants. To familiarize participants with the required movement speed, they received 24 and 12 practice trials for the first and second speed conditions, respectively. Practice trials were repeated only if participants were unable to perform any trial within the required speed range. In total, 30 trials were performed for each starting position.

The primary motor outcome measure was the onset timing difference between the shoulder extensor (i.e., m. posterior deltoid) and the elbow extensor (i.e., m. triceps lateral head). EMG signals were sampled at 1000 Hz, band-pass filtered (20–100 Hz), rectified, and normalized to the mean activity required to counter a 2 N·m torque (see: Pruszynski et al., 2008). EMG onset was defined as the time at which the mean EMG trace across all valid trials exceeded 3 SD above baseline (calculated as the mean muscle activity from −400 to −300 ms relative to movement onset) for at least 50 ms. Trials in which participants initiated their movement in the wrong direction were excluded from the analysis.

We chose this outcome measure because anticipatory muscle activation, which is crucial for movement stabilization, relies on the feedforward control mechanisms underlying the coordination of pure elbow movement (Bastian, 2006; Gribble & Ostry, 1999; Hollerbach & Flash, 1982). The cerebellum is crucial for the predictive control of intersegmental dynamics required for accurate temporal coordination between the shoulder and elbow muscles and for movement stabilization (Oh et al., 2025).

To measure the resilience of cerebellar motor function, we quantified the change in onset timing difference between shoulder and elbow muscle activity across the fast and slow conditions. When the control system is challenged by faster movement speeds, a shorter time interval between the anticipatory shoulder muscle activation relative to elbow activation would indicate a deterioration of anticipatory muscle timing (hence absence of resilience), which depends on the cerebellum.

#### **Motor timing task** (cerebellar specific outcome measure: variability in the internal timekeeping mechanism)

Motor timing ability was assessed using a rhythmic finger-tapping task performed on a computer keyboard. Participants were seated at a table and instructed to tap the spacebar in synchrony with a sequence of auditory tones presented at a fixed frequency. After 13 tones, participants continued tapping at the same pace without auditory cues for an additional 31 taps (**Figure 1B**).

The task was performed under three tempo conditions, defined by the inter-stimulus interval (ISI) between tones: (1) 600 ms (1.67 Hz), (2) 1,000 ms (1 Hz), and (3) 1,400 ms (0.71 Hz). The 1.67 Hz condition corresponds to the paradigm described previously by de Witte et al. (2026a, 2026b). Increasing the ISI was considered the task-specific stressor, as longer intervals between taps are known to increase variability and difficulty (Grube et al., 2010). For each condition, one practice trial was completed, followed by three test trials. The order of conditions was randomized across participants.

The primary outcome measure was the clock variance, computed based on the model of Wing and Kristofferson (1973), which reflects the variability in the internal timekeeping mechanism (see de Witte et al., 2026b for more details on the calculation of the outcome parameter). For each trial, inter-tap intervals were derived from the final 30 unguided taps. Intervals shorter than 150 ms (i.e., indicating a spacebar fault during tapping) or longer than twice the target ISI (i.e., indicating loss of attention) were removed. If an interval twice as long as the target interval was detected in the middle of a trial, the entire trial was excluded. We then calculated the mean clock variance across all valid trials for each condition.

Clock variance is considered a cerebellar-specific index of timing precision, as it reflects fluctuations in the internal timing mechanism rather than movement execution per se. Previous studies have shown that individuals with cerebellar lesions exhibit increased clock variance, indicating impaired predictive timing control mediated by cerebellar circuits (Ivry & Keele, 1989; Harrington, 2003; Franz et al., 1996).

To assess the resilience of this cerebellar-specific timing function, we quantified the change in clock variance between the most challenging (0.71 Hz) and the least challenging (1.67 Hz) tempo conditions. Under heightened task demands, no or only small increases in clock variance across tempo conditions would indicate greater resilience of the cerebellar-dependent timing mechanism.

#### **Postural stability task** (general sensorimotor outcome measure: postural sway)

Postural stability was assessed while participants maintained a quiet stance on a portable force plate (BTracks Inc.) with their hands placed on their hips and their gaze fixed on a cross 1-2 m in front of them.

The task was performed in four stance variations: (1) bipodal stance with eyes open (BSEO), (2) bipodal stance with eyes closed (BSEC), (3) semi-tandem stance with eyes open (TSEO), and (4) semi-tandem stance with eyes closed (TSEC) (**Figure 1C**). Participants from the oldest age group did not perform the latter condition (TSEC) for safety reasons. Changing the foot position from bipodal to semi-tandem and the elimination of visual feedback are considered as task-specific stressors, as both manipulations increase the postural control demands. The semi-tandem stance reduces the base of support and introduces asymmetry in weight distribution, thereby limiting the ability to make effective balance corrections and resulting in lower stability compared to the bipodal stance (Magalhães et al., 2022; Malcolm et al., 2021). Similarly, performing the task with eyes closed removes visual input, forcing greater reliance on proprioceptive and vestibular information, which are less precise for maintaining upright posture. Together, these conditions make the task progressively more challenging and sensitive to individual differences in balance control. For each condition, one familiarization trial was performed, followed by three test trials of 35 seconds. The order of the conditions was consistent across all participants (1: BSEO, 2:BSEC, 3:TSEO, 4:TSEC).

Postural sway was quantified using the prediction ellipse area, which represents the 95% confidence area enclosing the center-of-pressure trajectory in the anterior–posterior and medial–lateral directions. The prediction ellipse area provides a two-dimensional measure of the spatial variability of center-of-pressure displacement, with larger areas indicating greater postural instability. For each trial, the center-of-pressure time series from the final 30 seconds, in both axes, was used to compute the covariance matrix of sway dispersion. The orientation and magnitude of the principal axes of the resulting ellipse were derived from the eigenvalues and eigenvectors of this covariance matrix. Based on these parameters, the area of the 95% prediction ellipse was calculated according to the procedure described by Schubert & Kirchner (2013), which provides a standardized way to convert the covariance of the center-of-pressure signal into a 95% prediction ellipse representing the expected sway region.

The prediction ellipse area was used as an indicator of general sensorimotor function. Postural stability depends on the integration of visual, vestibular, and somatosensory information with motor output to maintain balance. Although the cerebellum also plays a key coordinating role to maintain postural stability, the prediction ellipse area reflects the overall spatial variability of center-of-pressure adjustments, which captures the combined contribution of multiple sensory and motor systems rather than cerebellar function alone. Therefore, this outcome measure represents a broader measure of general sensorimotor performance.

Resilience of this general sensorimotor measure was defined as the difference in performance between the most challenging (TSEC: for young and older adults; and TSEO: for the older-old adults) and the least challenging (BSEO) condition. Greater resilience of the sensorimotor system is reflected in no or smaller increases in prediction ellipse area across the conditions of standing balance.

### 2.4. Cognitive tasks

#### **Mental rotation task** (cerebellar specific outcome measure: reaction time)

The mental rotation task assessed the ability to mentally transform visual stimuli. Participants were seated at a table and completed the task on a laptop computer. All task conditions reported here were identical to those used in the study by de Witte et al. (2026a, 2026b), but are included to address a different research question in the present analysis. On each trial, a single letter (F, G, J, or R; Helvetica font) was presented in white on a black background. Participants were instructed to judge as quickly and accurately as possible whether the letter was presented in its standard (e.g., “R”) or mirror-reflected form (e.g., “Я”), by pressing the left arrow key with their left index finger or the right arrow key with their right index finger, respectively. The stimulus remained on the screen until a response was made, after which feedback appeared for 1 s (“correct” in green or “incorrect” in red) (**Figure 1D**). Stimuli were presented in random order, with an equal number of standard and mirror-reflected stimuli. After five practice trials, participants completed 144 experimental trials, separated by a 2 s inter-trial interval.

Task difficulty was determined by the degree of rotation at which the letter was presented. It could appear in seven different orientations: 0° (baseline), −15°, 15°, −75°, 75°, −135°, or 135°. Increasing the rotation angle was considered the task-specific stressor, as greater angular rotation requires more extensive mental transformation to align the rotated image with the upright orientation. For each orientation, 18 trials were presented, except for the baseline condition, which was presented in 36 trials.

To assess cognitive performance, we calculated the mean reaction time (RT) for each rotation magnitude, combining clockwise and counterclockwise angles of equal magnitude. Only correct trials were considered, to ensure that reaction times reflected the ability to mentally transform the stimuli rather than response errors.

This outcome measure can be considered a proxy of cerebellar-dependent cognitive function, as the cerebellum contributes to continuous internal transformations of spatial representations. Evidence from neuroimaging studies and studies in patients with cerebellar lesions indicates that cerebellar damage leads to slower or less precise mental rotation performance, when rotating objects or body parts (McDougle et al., 2022; Picazio et al., 2013).

Specifically, to assess the resilience of this cerebellar-specific cognitive function, we calculated the slope of the regression line fitted to the mean reaction times across all orientations (0°, 15°, 75°, and 135°). The slope quantified the increase in response time per degree of rotation, reflecting the efficiency of mental transformation. A flatter slope indicates less performance deterioration under higher task demands and, therefore, greater resilience of cerebellar-dependent cognitive processing.

#### **Spatial working memory task** (general cognitive outcome measure: percentage of fault answers)

The spatial working memory task assessed how well participants could encode, maintain, and recall the spatial positions of multiple targets presented for 2 s in a circular grid. After a further 2 s delay, a question mark appeared for 2 s at or adjacent to one of the target locations. Participants had to indicate whether a target had previously been presented at the location of the question mark by reaching to the “yes” or “no” button (**Figure 1E**). Participants were seated in front of a KINARM end-point robot and used the handle as a cursor to indicate their response on the screen. Feedback after each trial was provided in green (i.e., correct answer) or red (i.e., incorrect answer).

The number of targets varied from three to six, with higher numbers increasing task difficulty. This manipulation served as the task-specific stressor because maintaining and accurately recalling a larger number of spatial locations imposes greater demands on working memory. Specifically, increasing the number of targets raises memory load, heightens interference between locations, and increases attentional and updating demands, thereby making correct performance progressively more challenging. A total of 12 trials per difficulty level were presented, with trials from all difficulty levels intermixed in a randomized order.

The percentage of correct responses was calculated for each difficulty level. Successful performance requires the coordinated engagement of multiple neural systems supporting attention, encoding, maintenance, and retrieval of spatial information. Cerebellar circuits are known to support visuospatial working memory. However, task performance is also influenced by cortical regions such as the prefrontal and parietal cortices, which play dominant roles in maintaining and manipulating spatial information. Therefore, although cerebellar function may contribute to performance on this task, the outcome measure is best interpreted as a proxy for general cognitive capacity.

To assess the resilience of general cognitive function, we calculated the performance difference between the most challenging and the two least challenging conditions of the spatial working memory task. Specifically, we computed the mean percentage of correct responses for the two most challenging conditions (five and six targets) and subtracted from it the mean percentage of correct responses for the two least challenging conditions (three and four targets). Averaging across two conditions within each difficulty level increased the number of trials contributing to each estimate, thereby improving the reliability of the measure. No or smaller differences between most challenging and least challenging conditions reflect a greater ability to maintain task performance under increased cognitive load and thus indicate higher resilience of general cognitive function.

### 2.5. Statistical analyses

All statistical analyses were conducted using MATLAB (MathWorks, Natick, MA, version 2024b). For each task, we examined the effects of age group and the task-specific stressor on performance. To this end, a two-way repeated-measures ANOVA, using the *ranova*-function, was performed with age group as a between-subjects factor and task-specific stressor level as a within-subjects factor. This analysis tested the main effects of age and task difficulty, as well as their interaction, which reflects potential age-related differences in task resilience. When significant main or interaction effects were observed, post hoc Tukey-Kramer comparisons were performed to identify pairwise differences between age groups and between levels of the task-specific stressor.

To further examine these age-related differences in resilience on the cerebellar-specific and general outcome measures, a one-way ANOVA, using the *anova1*-function, with age as between-subjects factor was performed on the resilience measure derived from each task. These analyses tested for an overall effect of age group on resilience, representing the interaction effects identified in the repeated-measures ANOVA.

Finally, to investigate potential shared mechanisms of resilience across tasks and domains, we performed robust *Spearman* correlation analyses using robustly standardized resilience outcome measures from all five tasks. To enable comparison of relative differences in resilience across tasks and individuals, resilience outcome measures were normalized using robust z-scores computed within each age group. For each variable, the robust z-score was defined as 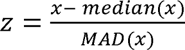, where *median(x)* represents the group-specific median and *MAD(x)* the unscaled median absolute deviation (i.e., *MAD(x) = median(|x – median(x)|*)). Correlations were computed separately for young and older adults. The older-old group was excluded from these analyses as the smaller sample size of the this group limited statistical power. All statistical tests were two-tailed, and significance thresholds were set at p < 0.05.

### 2.6. MRI acquisition

In addition to the two experimental sessions, including motor and cognitive assessments, one experimental session of ∼1 hour was conducted at the University Hospital of KU Leuven, Belgium. During this session Magnetic Resonance Images (MRI) were acquired from the whole brain, using a 3 Tesla Philips MRI Achieva dStream scanner equipped with a 32-channel receive-only head coil. A high-resolution three-dimensional T1-weighted structural image was acquired (repetition time (TR) = 9.7 ms; echo time (TE) = 4.6 ms; inversion time (TI) = 900 ms; flip angle = 8°; field of view (FOV) = 256 x 242 x 182 mm; voxel size = 0.89 x 0.89 x 1.0 mm; 182 sagittal slices).

The MRI dataset used in the present study was previously analyzed in de Witte et al. (2026a, 2026b). In those studies, cerebellar grey matter volume was quantified using anatomical parcellations (de Witte et al., 2026b) and combined anatomical and functional parcellation approaches (de Witte et al., 2026a). In the present study, the same dataset was used to investigate the structure-function relationship between cerebellar grey matter volumes based on a functional parcellation of the cerebellum (Nettekoven et al., 2024) and task-based resilience outcome measures.

All anatomical scans were first manually aligned to the anterior commissure to enhance segmentation precision, using SPM12 (Wellcome Trust Centre for Neuroimaging, London, UK). Subsequent voxel-based morphometry (VBM) preprocessing and analysis were performed with the Computational Anatomy Toolbox (CAT12.8.2) implemented in SPM12. Cerebellar region-of-interest (ROI) volumes were defined according to the asymmetric functional atlas introduced by Nettekoven et al. (2024), which delineates 32 cerebellar regions in MNI152 standard space. Before applying the atlas, it was spatially co-registered to the shooting template used by CAT12 to ensure consistent segmentation and normalization. Total grey matter volumes for each ROI, along with total intracranial volume (TIV), were automatically extracted during the VBM pipeline. Because hemispheric asymmetry was not examined in the current work, left and right homologous regions were combined, yielding 16 bilateral functional cerebellar ROIs: 4 motor, 3 action, 4 demand, and 5 sociolinguistic regions. For each functional domain, the total grey matter volume was computed as the sum of its constituent ROIs. To account for individual differences in head size, ROI-specific grey matter volumes were normalized by TIV prior to further analyses.

### 2.7. Predictive model

To assess the functional relevance of the cerebellar structural measures, we examined whether interindividual variability in grey matter volume within the four defined cerebellar functional regions (i.e. motor, action, demand and sociolinguistic) (King et al., 2019; Nettekoven et al., 2024) could predict resilience outcomes on motor and cognitive tasks. For this purpose, we employed a regularized linear regression model using ridge regression, which reduces overfitting by penalizing large regression coefficients and mitigates multicollinearity among correlated cerebellar region volumes. A fixed ridge penalty (λ = 1) was used to provide mild regularization while preserving interpretability of the model coefficients. Prior to model fitting, both cerebellar grey matter volumes and behavioral resilience outcomes were robustly standardized (z-scored) within each age group using the median and median absolute deviation (MAD) approach described earlier.

Model performance was quantified using the cross-validated coefficient of determination (R^2^), reflecting the proportion of variance in resilience outcomes that could be predicted from cerebellar grey matter volume. For each task and age group, R^2^ was computed using leave-one-out cross-validation. Specifically, for each participant, the ridge regression model was trained on data from all other participants, and the obtained model was then used to predict the left-out participant’s resilience score. More specifically, let *y_p_* denote the observed (z-scored) functional resilience outcome of a specific task for participant *p*, and 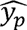 the corresponding cross-validated model prediction. The mean observed value across participants is denoted by 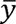 . The total sum of squares (*SS*_tot_) quantifies the total variance in the resilience outcome of the specific task: 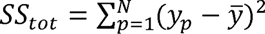. The residual sum of squares (*SS*_res_) captures the variance unexplained by the model 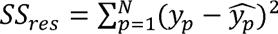. The coefficient of determination (R^2^) is then defined as 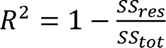.

The reported R^2^ value indicates how accurately grey matter volume across the four cerebellar functional regions predicts individual differences in functional resilience outcomes on the tasks. Positive R^2^ values indicate that the variability in cerebellar grey matter volume across the four functional regions explains a portion of the between-participant variance in functional task resilience outcomes. A R^2^ value of zero indicates that the model predicts individual outcomes no better than predicting the group average, whereas negative R^2^ values indicate that the model predicts individual outcomes worse than the mean-based prediction.

To contextualize model performance, R^2^ values were interpreted relative to the task-specific noise ceiling, representing the maximum explainable variance given the intrinsic reliability of each behavioral outcome. Noise ceilings were estimated using odd-even split-half correlations of resilience outcomes (z-scored) across all participants (young and older adults combined), providing an index of the internal consistency of the task measure (van Bree et al., 2025). Higher noise ceiling values indicate greater reliability of the task measure. Comparing model R^2^ values to the noise ceiling allow us to determine how much of the reliable variance in the resilience outcomes across tasks could be captured by cerebellar grey matter volume. This comparison allows for a more meaningful interpretation of model performance, especially across tasks that differ in their measurement reliability.

To assess the stability and significance of individual regression coefficients, ridge regression coefficients were additionally evaluated using bootstrap resampling (5,000 iterations). Percentile-based confidence intervals and two-sided bootstrap p-values were computed for each cerebellar region, with p-values corrected for multiple comparisons using the Benjamini-Hochberg false discovery rate (FDR) procedure (Benjamini & Hochberg, 1995). Analyses were restricted to young and older adult groups, as MRI data were only available for a subset of older-old adults (19 of the 30), limiting statistical power in that group.

## Results

We examined performance across three age groups on five tasks (three motor, two cognitive), each providing either a cerebellar-specific or a general outcome measure. Each task comprised multiple levels of difficulty defined by a task-specific stressor, enabling us to assess the resilience of these outcomes and how resilience varies with healthy aging.

### 3.1. Cerebellar-specific motor measures remain resilient across aging

We tested whether cerebellar motor function is maintained during healthy aging using two motor tasks from which we derived cerebellar-specific outcome measures. In the first task, anticipatory muscle activation was assessed using a pure elbow motion task in which the timing of the first activation burst of the shoulder muscle was measured relative to that of the elbow muscle (for a full description of the behavior in this task, Matthijs et al. (2025)). This anticipatory activation, a process that critically depends on cerebellar function, is required to counteract for the interaction torque generated by forearm rotation. Our results provided no evidence for age-related changes in this anticipatory shoulder–elbow activation timing (**Figure 2A**; main effect of age group: F(2,103) = 0.76, p = 0.47), indicating that all age groups were able to effectively counteract the interaction torque at the shoulder. The shoulder muscle was activated 13 ± 19 ms, 10 ± 16 ms, and 14 ± 20 ms before elbow muscle activation in young, older, and older-old adults, respectively.

**Figure 2:**
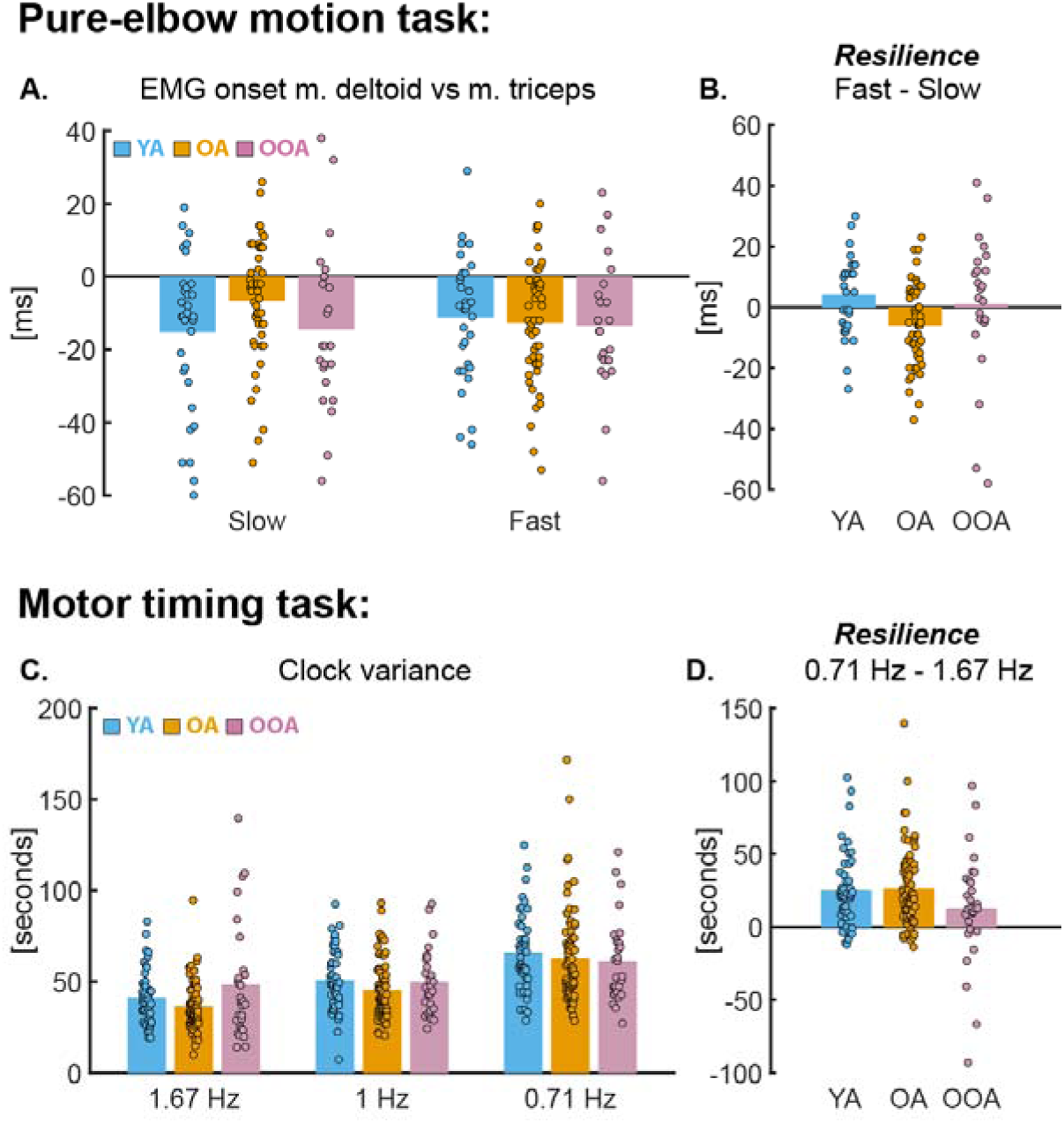
**A)** Timing of EMG onset of the shoulder muscle (m. deltoid) relative to the elbow muscle (m. triceps) during pure elbow extension movements. **B)** Difference in EMG onset timing between the fast (most challenging) and slow (least challenging) speed conditions, representing the resilience measure of the pure elbow motion task (cerebellar-specific outcome). Higher values indicate lower resilience (i.e., poorer task performance under increased demands). **C)** Clock variance reflecting timing variability during the motor timing task. **D)** Difference in clock variance between the 0.71 Hz condition (most challenging) and the 1.67 Hz condition (least challenging), representing the resilience measure of the motor timing task (cerebellar-specific outcome). Higher values indicate lower resilience (i.e., poorer task performance under increased demands). **All panels:** Each dot represents an individual participant. Bar height indicates the group mean for each condition and age group. Blue = young adults; orange = older adults; purple = older-old adults.

While age did not affect this anticipatory muscle timing, we further investigated whether task performance could be maintained under increased motor demands. Although increasing forearm movement speed placed greater demands on predictive motor coordination to compensate for the increased interaction torque at the shoulder, we found that the relative timing between activation of the shoulder and elbow muscles remained similar across the slow and fast speed conditions (**Figure 2A**; main effect of stressor: F(1,103) = 0.036, p = 0.85). The timing of shoulder muscle activation relative to elbow muscle activation was 12 ± 19 ms and 12 ± 17 ms in the slow and fast speed conditions, respectively. This suggests that anticipatory muscle timing between the shoulder and elbow muscles was not affected by higher task demands.

Interestingly, when evaluating the resilience of anticipatory muscle timing, older and older-old adults did not show greater performance declines under increased task demands than young adults. In fact, older adults exhibited earlier anticipatory activation of the shoulder relative to the elbow in the fast-speed condition (OA = 6 ± 23 ms earlier than in the slow-speed condition), whereas anticipation decreased in the young and older-old groups (YA = 4 ± 13 ms later; OOA = 1 ± 14 ms later), resulting in a significant interaction effect between age and task stressor (F(2,103) = 4.11, p = 0.019). Consequently, the cerebellar-specific outcome measure indicated better anticipatory muscle timing in older adults compared with young adults (YA vs. OA: p = 0.019, d = -0.75, 95% CI [-1.21, -0.29]). However, when considering only the fast-speed condition, the relative timing between shoulder and elbow muscle activation was nearly identical across age groups (YA = -12 ms; OA = -13 ms; OOA = -13 ms). No additional differences in resilience were observed between age groups (YA vs. OOA: p = 0.772, d = -0.16, 95% CI [-0.69, 0.37]; OA vs. OOA: p = 0.190, d = 0.41, 95% CI [-0.08, 0.90]) (**Figure 2B**).

In motor timing task, we measured clock variance as an index of cerebellar motor function, an index of variability in the internal timekeeping mechanism (results from the 1.67 Hz condition are presented in de Witte et al., 2026b). Consistent with the findings for anticipatory muscle timing, this cerebellar-specific measure also did not differ across age groups (**Figure 2C**; main effect of age group: F(2,154) = 2.40, p = 0.094). This suggests that the cerebellar-dependent mechanism responsible for generating consistent rhythmic intervals remains stable with age. Mean clock variance was 53 ± 11 ms in young adults, 48 ± 14 ms in older adults, and 53 ± 16 ms in oldest-old adults.

Regarding the effect of the task-specific stressor, increasing the inter-stimulus interval led to higher clock variance, indicating greater variability in the internal timing mechanism (**Figure 2C**; main effect of stressor: F(2,154) = 54.45, p < 0.001). Post hoc Tukey–Kramer comparisons confirmed that all task levels differed significantly from one another (all p < 0.001). Clock variance increased from 42 ± 19 ms at the 600 ms interval, to 49 ± 16 ms at the 1,000 ms interval, and to 63 ± 24 ms at the 1,400 ms interval, demonstrating that task performance deteriorated as task demands increased.

When assessing the resilience of the internal timekeeping mechanism, we again found no evidence of age-related differences in clock variance across increasing task demands (interaction effect between age group and stressor: F(4,154) = 1.62, p = 0.168). For all age groups, performance deteriorated in the most difficult condition compared with the easiest condition, and the magnitude of this deterioration was similar across groups: 24 ± 26 ms in young adults, 26 ± 26 ms in older adults, and 13 ± 39 ms in older-old adults (YA vs. OA: p = 0.967, d = 0.05, 95% CI [-0.31, 0.41]; YA vs. OOA: p = 0.188, d = -0.38, 95% CI [-0.85, 0.09]; OA vs. OOA: p = 0.089, d = -0.44, 95% CI [-0.87, -0.01]) (**Figure 2D**). If anything, the increase in clock variance was the smallest for the older-old adults.

Taken together, both cerebellar-specific motor tasks indicate that older and older-old adults maintained motor performance to a similar extent as young adults, even under increased task demands.

### 3.2. General sensorimotor measure shows age-related changes, with resilience declining already in older age

Having established that cerebellar-specific motor functions remain resilient with age, we next asked whether this resilience extends to motor abilities that rely more broadly on general sensorimotor processes. To test this, we examined age effects on a postural stability task, which engages multiple sensory and motor systems. Across all conditions of this quiet standing task, we found that postural sway increased with age (**Figure 3A**; main effect of age group: F(2,156) = 38.1, p < 0.001). Mean prediction ellipse area was 0.934 ± 0.486 m² in young adults, 1.244 ± 0.571 m² in older adults, and 2.274 ± 1.096 m² in oldest-old adults, with all age groups differing significantly (all p < 0.001). This indicates that the spatial variability of center-of-pressure adjustments, capturing general sensorimotor function, worsened with age.

**Figure 3:**
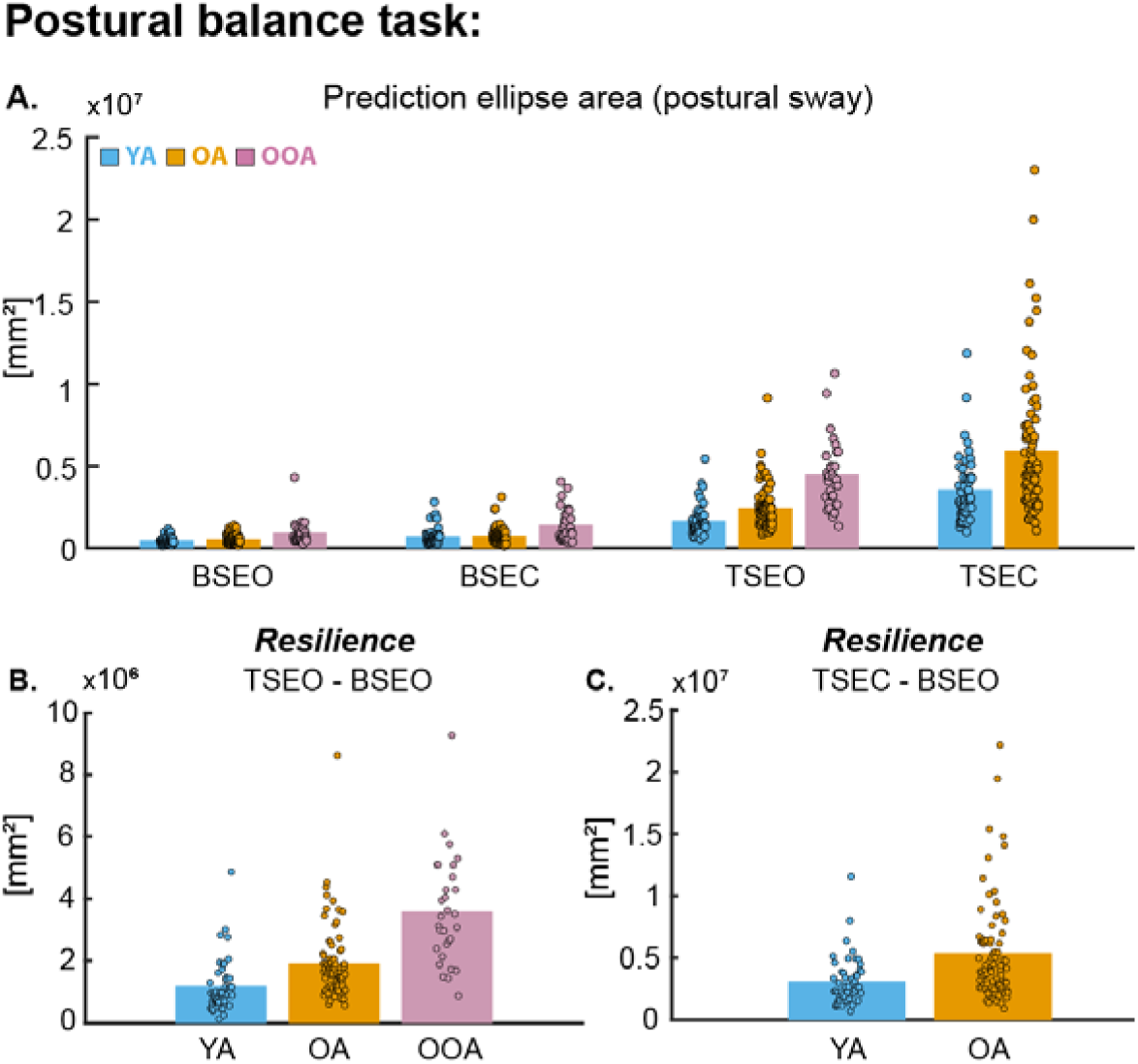
**A)** Postural sway during the postural balance task. **B)** Difference in postural sway between the tandem stance with eyes open (most challenging for older-old adults) and the bipodal stance with eyes open (least challenging), representing the resilience measure of the postural balance task (general sensorimotor outcome). Higher values indicate lower resilience (i.e., poorer task performance under increased demands). **C)** Difference in postural sway between the tandem stance with eyes closed (most challenging for older adults) and the bipodal stance with eyes open (least challenging). Higher values indicate lower resilience (i.e., poorer task performance under increased demands). **All panels:** Each dot represents an individual participant. Bar height indicates the group mean for each condition and age group. Blue = young adults; orange = older adults; purple = older-old adults.

The task-specific stressors also had a strong effect on postural stability: closing the eyes or narrowing the base of support produced progressively poorer performance (**Figure 3A**; main effect of stressor: F(2,156) = 367.78, p < 0.001). Across age groups, mean postural sway was 0.639 ± 0.427 m² in the bipodal stance eyes open condition, 0.953 ± 0.670 m² in the bipodal stance and eye closed condition, and 2.860 ± 1.697 m² in the tandem stand and eyes open condition. Each increment in task difficulty significantly increased postural sway (all p _≤_ 0.001), confirming that more challenging conditions resulted in poorer stability. Young and older participants additionally performed a semi-tandem stance with eyes closed, which further amplified postural sway to 4.719 ± 3.662 m², a significant increase relative to the tandem stance and eyes open condition (p < 0.001). This highlights how largely stability deteriorates when reduced sensory information is combined with a constrained base of support.

Most notably, when assessing the resilience of general sensorimotor function across age groups, we observed that postural stability declined more in the older groups as task demands increased (interaction effect between age group (YA, OA and OOA) and stressor (3 levels): F(4,156) = 31.19, p < 0.001). In addition, the difference in mean prediction ellipse area between the tandem stance and eyes open condition and the bipodal stance and eyes open condition (resilience outcome for this task) worsened with age (main effect of age group (F(2,156) = 36.71, p < 0.001), with all groups differing from each other (**Figure 3B**: YA vs. OA: p = 0.002, d = 0.70, 95% CI [0.33, 1.06]; YA vs. OOA: p < 0.001, d = 1.91, 95% CI [1.37, 2.45]; OA vs. OOA: p < 0.001, d = 1.22, 95% CI [0.77, 1.67]). The increase in postural sway from bipodal stance and eyes open condition to tandem stance and eyes open condition was 1.174 ± 0.830 m² in young adults, 1.914 ± 1.181 m² in older adults, and 3.576 ± 1.757 m² in older-old adults, demonstrating a clear age-related decline in general sensorimotor resilience. Consistent with this pattern, the increase in sway from bipodal stance and eyes open condition to tandem stance and eyes closed condition was substantially larger in older adults (5.363 ± 4.050 m²) than in young adults (3.083 ± 1.947 m²) (main effect of age (YA vs OA): F(1,127) = 14.94, p < 0.001; **Figure 3C**). This further highlights reduced resilience in a general sensorimotor task with advancing age.

### 3.3. Cerebellar-specific cognitive measure shows age-related changes, with resilience declining in older-old age

Given the cerebellum’s role beyond motor control, we next examined whether comparable resilience patterns emerged in the cognitive domain. To this end, reaction time was assessed using a mental rotation task, providing a cerebellar-specific cognitive measure (results from this task are also presented in de Witte et al., 2026b). Reaction time increased with age (main effect of age group: F(2,157) = 77.43, p < 0.001). Furthermore, the degree of letter rotation, which served as the task-specific stressor, also led to a significant increase in reaction time (main effect of stressor: F(3,157) = 420.35, p < 0.001). Mean reaction times were 1.08 ± 0.34 s for 0°, 1.12 ± 0.35 s for 15°, 1.29 ± 0.38 s for 75°, and 1.53 ± 0.45 s for 135° rotation. Post hoc Tukey–Kramer comparisons showed that all rotation levels differed significantly from each other (all p < 0.001). These results indicate that greater rotation angles increased task difficulty, as reflected in longer reaction times required for mental transformation of the letters (**Figure 4A**).

**Figure 4:**
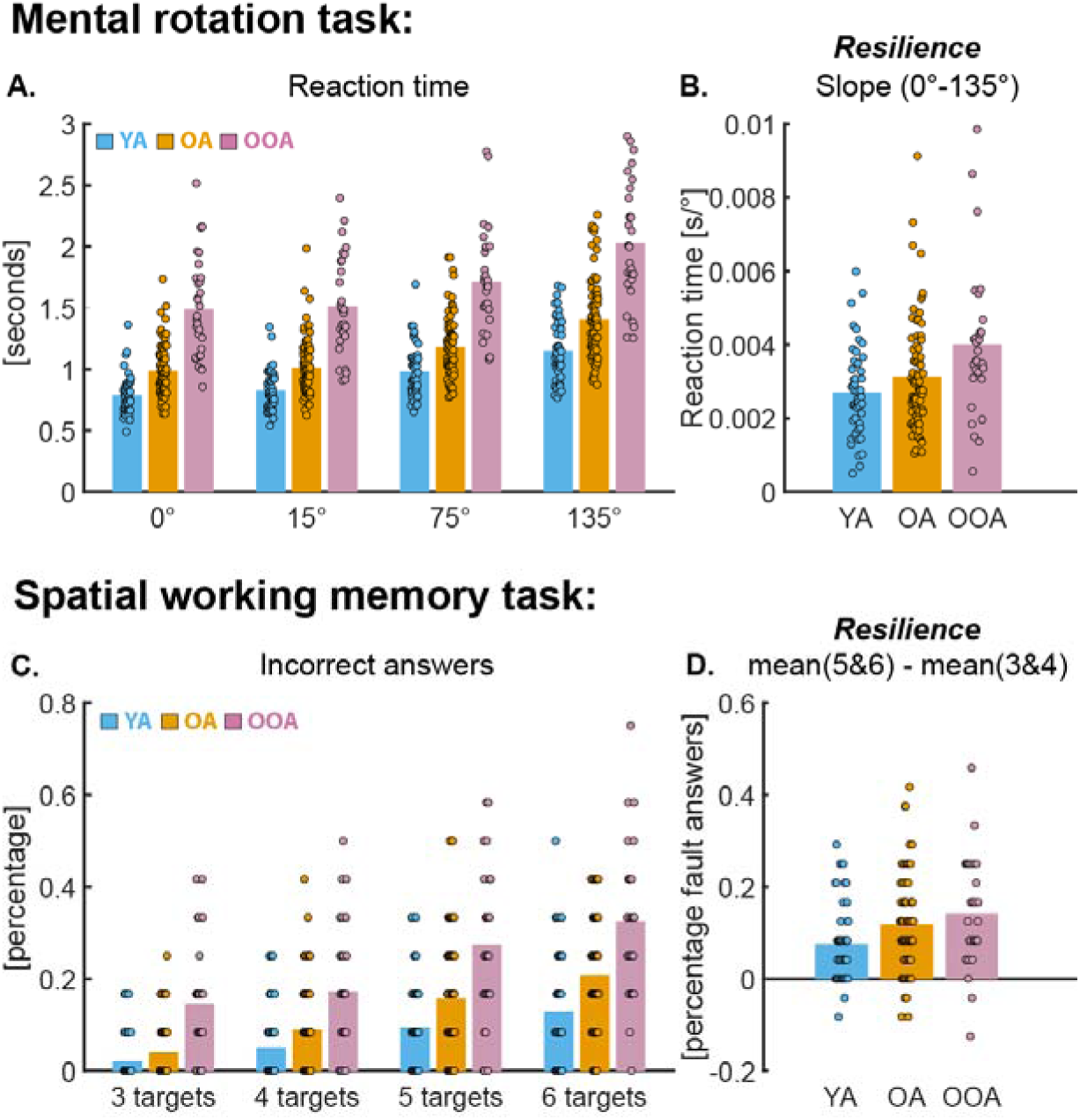
**A)** Reaction time during the mental rotation task. **B)** Slope of reaction times across all rotation conditions, representing the resilience measure for the mental rotation task (cerebellar-specific outcome). Higher values indicate lower resilience (i.e., poorer task performance under increased demands). **C)** Percentage of incorrect responses during the spatial working memory task. **D)** Difference between the mean number of incorrect responses in the most demanding conditions (5–6 targets) and the least demanding conditions (3–4 targets), representing the resilience measure for the spatial working memory task (general cognitive outcome). Higher values indicate lower resilience (i.e., poorer task performance under increased demands). **All panels:** Each dot represents an individual participant. Bar height indicates the group mean for each condition and age group. Blue = young adults; orange = older adults; purple = older-old adults.

Interestingly, the effect of increasing the rotation angle on reaction time was more pronounced with age (interaction effect between age group and stressor: F(6,157) = 5.36, p < 0.001). To assess resilience more directly, we examined the reaction-time slope across all rotation angles, as this metric provides the most sensitive cerebellar-specific outcome. A one-way ANOVA revealed that the reaction-time slope became steeper with age (**Figure 4B**; main effect of age group: F(2,157) = 7.07, p = 0.001). Post hoc comparisons confirmed that the increase in reaction-time cost under higher task demands was significantly greater in older-old adults than in both young (YA vs. OOA: p < 0.001, d = 0.85, 95% CI [0.38, 1.32]) and older adults (OA vs. OOA: p = 0.019, d = 0.53, 95% CI [0.10, 0.95]), whereas young and older adults did not differ significantly (YA vs. OA: p = 0.250, d = 0.31, 95% CI [-0.05, 0.66]). Overall, these findings indicate that the ability to mentally transform a visual stimuli, which relies on internal cerebellar processes, declines gradually with advancing age and becomes most apparent after 80 years of age.

### 3.4. General cognitive measure shows age-related changes, with resilience declining already in older age

Having identified that the cerebellar-specific cognitive measure showed decreased resilience emerging in the older-old group, we next investigated whether a similar pattern would be observed in a working memory task, which relies more heavily on general cognitive processing abilities. Performance on this task showed a significant age-related decline, as indicated by an increase in the percentage of incorrect responses with age (main effect of age group: F(2,157) = 43.87, p < 0.001). Furthermore, increasing the number of targets, which imposed greater demands on working memory, led to a significant increase in response errors (**Figure 4C**; main effect of stressor: F(1,157) = 168.76, p < 0.001), indicating that remembering target locations became increasingly difficult as task demands increased.

When evaluating how participants coped with the increasing task demands, we observed that resilience of working memory declines with age (**Figure 4D**; interaction effect between age group and stressor: F(2,157) = 4.62, p = 0.011). The older-old adults made more errors than young adults at higher difficulty levels (Post hoc Tukey: YA vs. OOA: p = 0.013, d = 0.55, 95% CI [0.09, 1.01]), whereas differences between young and older adults (YA vs. OA: p = 0.051, d = 0.51, 95% CI [0.15, 0.87]) and between older and older-old adults (OA vs. OOA: p = 0.522, d = 0.09, 95% CI [-0.33, 0.51]) were not significant. Overall, these findings indicate that resilience in this general cognitive measure declines with advancing age.

### 3.5. Resilience is domain- and task-specific and does not generalize across cerebellar-specific or general motor and cognitive measures

The previous analyses showed distinct age-related patterns of resilience across the five tasks. In the motor domain, resilience in the cerebellar-specific outcome measures was preserved across the lifespan. Effect sizes showed no meaningful differences between young adults and either older group, and even suggested a slight advantage in older adults. In contrast, resilience in the general sensorimotor measure declined progressively with age, with medium effects in older adults and large effects in older-old adults, indicating substantial vulnerability to increasing task demands (**Figure 5**).

**Figure 5:**
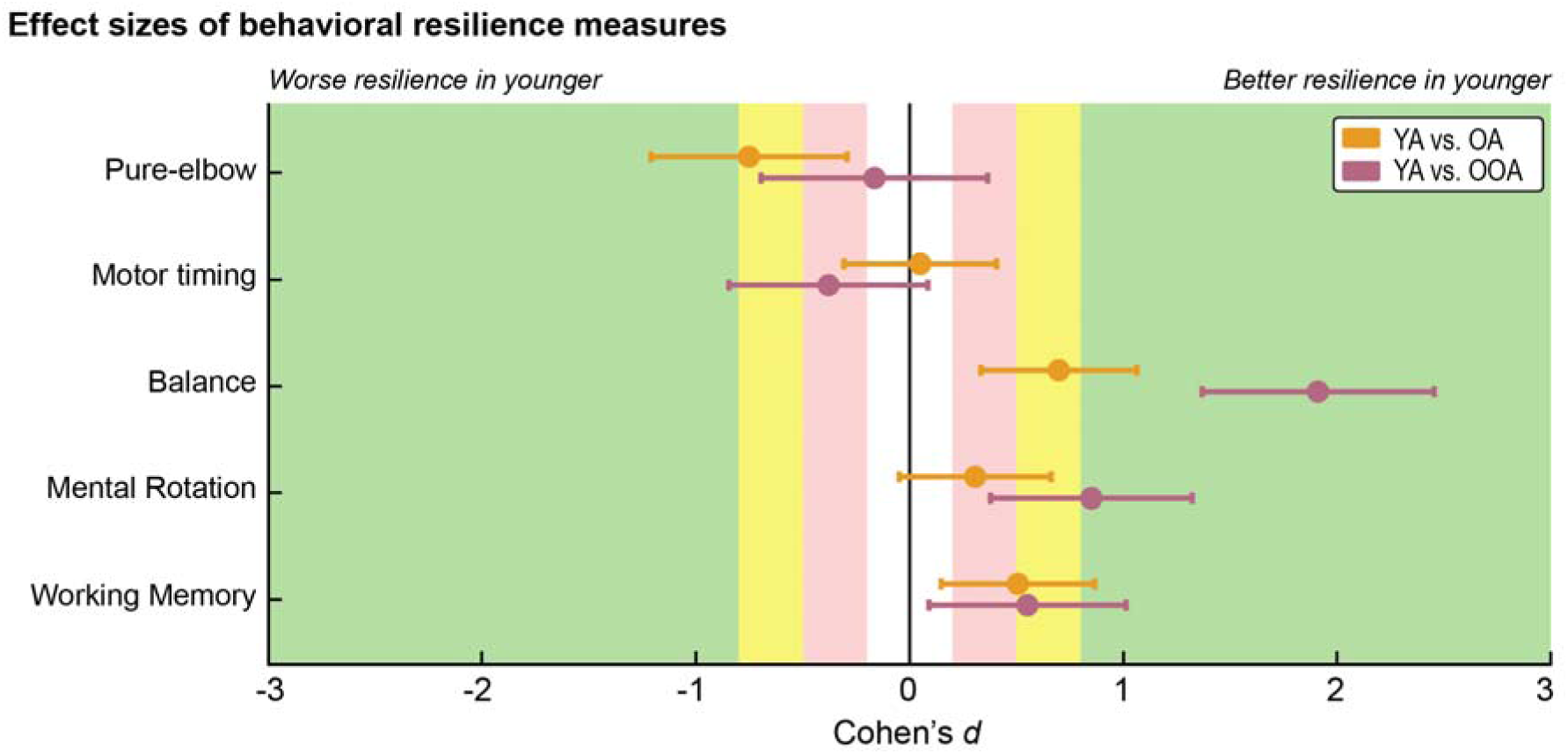
Overview of the effect sizes across all resilience outcomes of the different tasks. Each dot represents the effect size form statistical comparisons (one-way ANOVA) between two age groups (orange: young vs. older adults; purple: young vs. older-old adults). Horizontal error bars indicate 95% confidence intervals. Negative values reflect worse resilience outcome measures in the younger age group, while positive values reflect better resilience outcome measures in the younger age group. Cohen’s d effect sizes could be interpreted as follows: <0.2: no effect (white); 0.2-0.5: small effect (red); 0.5-0.8: medium effect (yellow); >0.8: large effect (green).

In the cognitive domain, a more graded pattern emerged. For the cerebellar-specific cognitive measure, increasing task demands had only a small effect in older adults compared with young adults, suggesting relative preservation of resilience. However, this advantage disappeared in the older-old group, where large effects indicated marked decline. By contrast, the general cognitive measure already showed a medium reduction in resilience in older adults, which increased slightly further in older-old adults (**Figure 5**). Together, the effect sizes across task outcomes suggest that cerebellar-specific functions are relatively more resilient than general functions in both domains, although this advantage appears to diminish for cognitive processes in advanced age.

Having established these patterns of resilience across the motor and cognitive domains, we next asked whether resilience reflects a general person-level characteristic, such that individuals who are more resilient in one task would also be more resilient in other tasks, rather than a task-specific phenomenon. If so, resilience measures should correlate across tasks. To evaluate this possibility, we computed pairwise correlations between all resilience outcomes (z-scores), separately for young and older adults. However, in neither age group was greater resilience on one task associated with greater resilience on any other task (all p > 0.05). Across all comparisons, correlation coefficients remained below r = 0.30, indicating that even the largest associations were small to medium in size and did not provide meaningful evidence that resilience reflects a shared person-level trait across domains (**Figure 6**).

**Figure 6:**
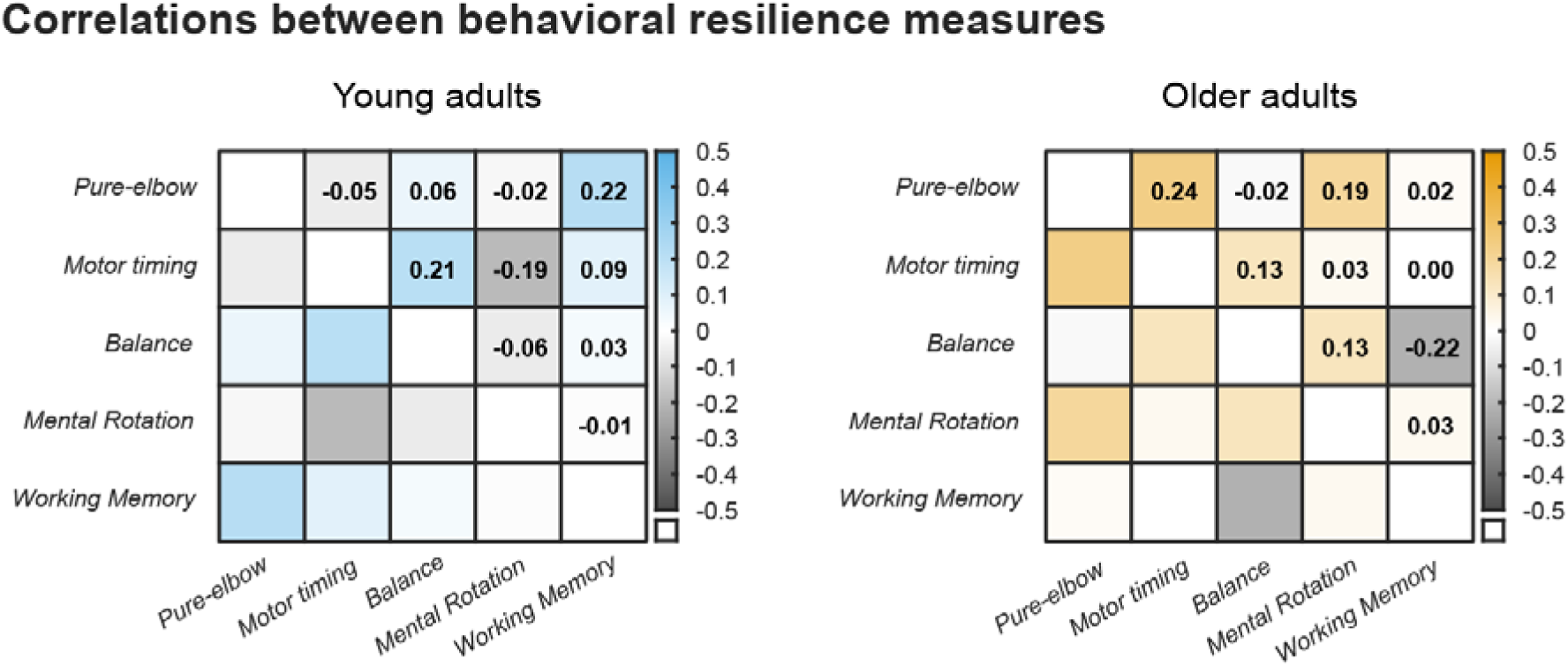
Overview of the correlations between resilience measures across tasks for young adults (blue) and older adults (orange). Color intensity indicates the strength and direction of the associations: darker hues of blue or orange represent more positive correlations, whereas grey tones indicate more negative correlations. Correlation coefficients are displayed in the upper triangle.

The absence of association between the cerebellar-specific motor measures suggests that preserved resilience in one cerebellar-dependent motor function does not necessarily predict resilience in another. Likewise, resilience in general sensorimotor and cognitive functions appeared independent, and no cross-domain relationships emerged between motor and cognitive resilience. Together, these findings suggest that resilience to increasing task demands is largely domain- and task-specific rather than reflecting a generalized personal characteristic. In the context of cerebellar reserve, this indicates that preserved cerebellar-specific resilience with aging may not directly translate into enhanced resilience in other motor or cognitive functions.

### 3.6. Cerebellar grey matter volume does not predict resilience in motor or cognitive measures

Finally, having demonstrated strong resilience in cerebellar-specific outcome measures during healthy aging, we examined whether this functional resilience was associated with age-related deterioration of cerebellar structure. Whole-brain MRI data, previously reported in de Witte et al. (2026a, 2026b), were analyzed here in relation to resilience outcomes to investigate potential associations with cerebellar grey matter volume. Using ridge linear regression, we found that interindividual differences in grey matter volume within the four predefined cerebellar functional regions (i.e., motor, action, demand, and sociolinguistic) could not predict resilience outcomes for the cerebellar-specific measures. Likewise, cerebellar grey matter volume also failed to predict resilience in general sensorimotor or cognitive measures.

Across all models, the explained variance (R²) was negative, indicating that model predictions based on cerebellar grey matter volume performed worse than a simple mean-based prediction model (**Figure 7**; pure elbow motion task: R^2^_YA_= -0.41 ± 0.25, R^2^_0A_= -0.15 ± 0.11; motor timing task: R^2^_YA_= -0.32 ± 0.13, R^2^_0A_= -0.15±0.08; postural balance task R^2^_YA_= -0.29 ± 0.14, R^2^_0A_= -0.25 ± 0.10: mental rotation task: R^2^_YA_ = -0.34 ± 0.15, R^2^_0A_= -0.14 ± 0.08; working memory task: R^2^_YA_= -0.21 ± 0.14, R^2^_0A_= -0.06 ± 0.08). Consistent with this, none of the models approached the respective noise ceiling values, reinforcing the absence of a meaningful structure-function relationship.

**Figure 7:**
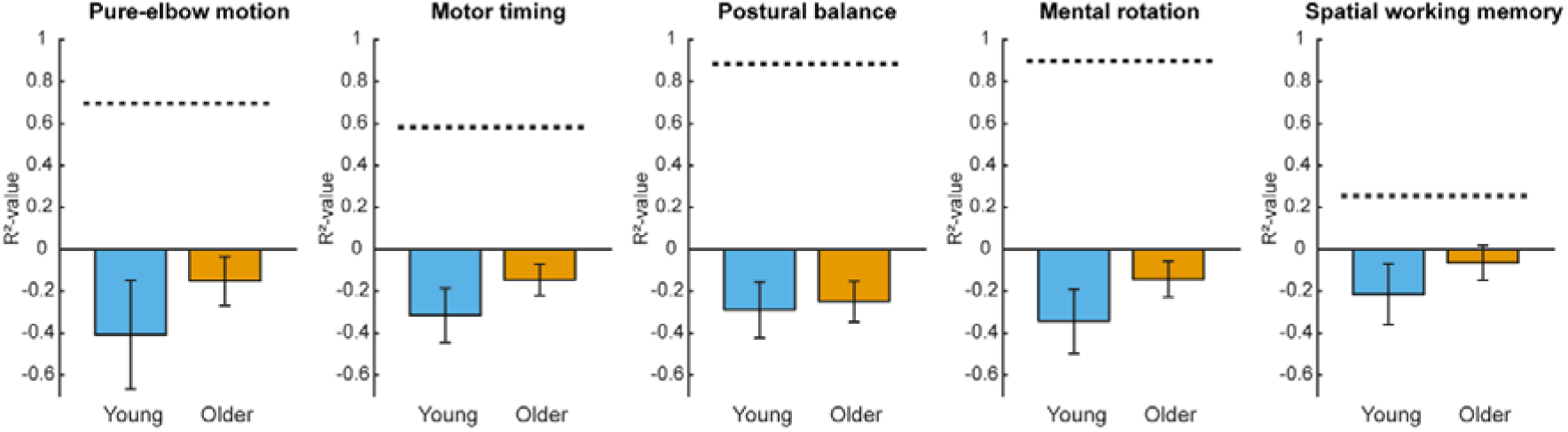
For each task, the R² value is shown for young (blue) and older (orange) adults, indicating the proportion of variance in the resilience outcome explained by grey matter volume across the four cerebellar functional regions. The dotted horizontal line represents the noise ceiling for each task, reflecting the maximum variance that could be explained by the predictive model.

## Discussion

### 4.1. Summary

In the present study, we investigated how resilience to increasing task demands varied across cerebellar-specific and general functional outcomes and across motor and cognitive domains in healthy aging. Our results demonstrated that, in the motor domain, cerebellar-specific motor outcome measures remained functionally preserved with advancing age, showing resilience under increased task demands even in adults over 80 years of age. In contrast, general sensorimotor outcome measures exhibited pronounced age-related declines and reduced resilience. In the cognitive domain, both cerebellar-specific and general outcome measures showed age-related reductions in resilience. However, cerebellar-specific cognitive processes appeared somewhat more resilient in older adults, with clearer declines emerging only in the older-old age group. Interestingly, resilience did not generalize across tasks: resilience measures were uncorrelated across tasks, even among those relying on cerebellar motor mechanisms. Moreover, resilience in any of the motor or cognitive outcomes was not predicted by cerebellar grey matter volume, indicating that resilience was not supported by corresponding structural preservation.

### 4.2. Preserved cerebellar-specific motor resilience across aging supports the cerebellar motor reserve hypothesis

Our present findings provide converging evidence for a cerebellar motor reserve hypothesis, which proposes that motor function can remain resilient to age-related cerebellar degeneration due to a reserve capacity within the cerebellum (Elbaz et al., 2013; Fleischman et al., 2015). Specifically, cerebellar-dependent motor mechanisms, as assessed by feedforward control during the pure elbow motion task and internal timing during the motor timing task, were functionally maintained across the adult lifespan, even under increased task demands. Importantly, inter-individual differences in resilience in performance on these cerebellar-specific motor tasks were not predicted by structural measures of cerebellar grey matter volume within the predefined functional parcellation of the cerebellum (Nettekoven et al., 2024). This dissociation between maintenance of motor function and age-related structural decline suggests that the cerebellum can act as a motor reserve, protecting cerebellar motor function from age-related cerebellar structural degeneration.

Support for this cerebellar motor reserve hypothesis comes from a broader body of experimental work demonstrating maintained performance in tasks that are strongly dependent on cerebellar motor processing in older adults. Across multiple studies, no age-related declines have been reported in motor tasks such as grip-load force coupling, inter-joint coordination, motor timing, sensory attenuation, and motor adaptation, which involve processes widely considered markers of cerebellar motor control (Duchek et al., 1994; Gilles & Wing, 2003; Hermans et al., 2025; Huang & Ahmed, 2014; Lee et al., 2007; Parthasharathy et al., 2022; Vandevoorde & Orban de Xivry, 2019; Wolpe et al., 2016). Although these studies did not directly relate functional performance to cerebellar structure, longitudinal neuroimaging work has consistently demonstrated cerebellar volume loss during healthy aging (Raz et al., 2005). Notably, recent work from our research group (de Witte et al., 2026b), conducted in the same cohort as in the present study, directly examined both functional and structural cerebellar outcomes across seven motor tasks, including the two cerebellar-specific tasks reported here. While cerebellar-dependent functional performance remained preserved across all tasks, clear age-related reductions in total cerebellar grey and white matter volume were observed, providing direct evidence for a dissociation between cerebellar structure and motor function. In addition, in a second study, our group showed that cerebellar grey matter volume could not account for inter-individual variability in cerebellar motor function, thereby indicating a lack of structure-function relationship consistent with the cerebellar reserve hypothesis (de Witte et al., 2026a). Collectively, these findings support the notion that the cerebellum can act as a motor reserve, maintaining motor performance despite substantial age-related structural degeneration.

At the same time, other studies have reported associations between cerebellar atrophy and motor performance deficits in aging (Bernard & Seidler, 2014; Hogan, 2004), lending support to a structural hypothesis in which age-related cerebellar degeneration directly contributes to motor decline. However, these studies typically rely on measures that do not specifically isolate cerebellar-dependent processes. As a result, the reported associations may reflect contributions from multiple neural systems rather than cerebellar function per se. Although this perspective contrasts with the present findings supporting a cerebellar motor reserve, the two views are not mutually exclusive. If the cerebellum indeed acts as a motor reserve, its protective capacity is likely finite, implying the existence of a threshold or “tipping point” beyond which further structural degeneration results in functional decline. Within this framework, cerebellar structure would become a limiting factor for motor performance only after reserve capacity has been sufficiently depleted. However, based on our data and prior evidence, it appears unlikely that this tipping point is reached during healthy aging, at least up to the ninth decade of life, as cerebellar motor performance remained preserved even in advanced age despite pronounced structural decline (de Witte et al., 2026b). Collectively, these findings suggest that while a minimum level of cerebellar structural integrity is necessary for motor function, age-related cerebellar degeneration within a healthy range does not appear to constrain cerebellar-specific motor outcomes.

### 4.3. Cerebellar-specific resilience appears more robust in the motor than in the cognitive domain

Although the cerebellum contributes to a wide range of functions beyond motor control (Massara et al., 2025; Rudolph et al., 2023; Schmahmann, 2019), the present findings suggest that cerebellar reserve capacity may be expressed more strongly in the motor domain than in cerebellar-dependent cognitive processes. Specifically, our findings showed that cerebellar-specific motor outcome measures remained preserved across the lifespan, whereas cerebellar-specific cognitive outcomes showed a modest decline in older adults and a more pronounced age-related decline in the older-old adults (≥80 years), despite widespread age-related structural decline of the cerebellum. This pattern raises the question of why cerebellar reserve appears to support resilience in motor-related functions better than other cerebellar-specific operations such as mental rotation.

One possible explanation for the stronger resilience of motor-related cerebellar processes could have been that reserve capacity supporting cerebellar-dependent cognitive processes is depleted earlier in the lifespan due to a higher rate of age-related degeneration in cerebellar regions that support cognitive functions. Previous work has indeed suggested that age-related structural changes in the cerebellum are spatially heterogeneous, with posterior regions implicated in cognitive processing exhibiting more pronounced age-related degeneration than regions primarily involved in motor control (Uquillas et al., 2024; Wang et al., 2024). However, analyses based on both the present dataset and on a subset of age-matched participants of the Cam-CAN (Cambridge Centre of Ageing and Neuroscience repository) cross-sectional population-based lifespan dataset (Shafto et al., 2014; Taylor et al., 2017) indicate that cerebellar grey matter volume declines at comparable rates across anterior and posterior regions, as well as across the four functional regions that can be defined in the cerebellum (de Witte et al., 2026a). These findings suggest that age-related changes in cerebellar structure cannot explain the observed differences in resilience outcomes between motor and cognitive functions. Consistent with this interpretation, grey matter volume within any of the four predefined cerebellar functional regions (King et al., 2019; Nettekoven et al., 2024) failed to predict resilience in the cerebellar-dependent cognitive task, indicating that cerebellar structure alone is insufficient to explain cognitive outcomes associated with cerebellar processing.

Another explanation may lie in differences in the organization of cortico–cerebellar connectivity. Similar to the cerebral cortex, where association areas such as the prefrontal and parietal cortices integrate input from multiple cortical regions to a greater extent than primary sensory or motor areas (Bertolero et al., 2015; Yeo et al., 2014, 2015), cerebellar regions may differ in the degree to which they receive convergent cortical input. Although relatively little is known about whether each cerebellar region receives input from a restricted cortical area or integrates signals from multiple cortical networks, recent work examining cortico–cerebellar connectivity has shown that cerebellar regions associated with higher-order cognitive functions exhibit greater degrees of cortical convergence than motor-related cerebellar regions (King et al., 2023). Importantly, prefrontal association areas, which show strong functional coupling with cerebellar regions supporting cognitive functions (King et al., 2023), are known to be particularly vulnerable to age-related structural decline compared with more posterior cortical areas (Lemaitre et al., 2012; Raz et al., 1997). Consequently, it is possible that greater convergence, and especially the strong functional coupling with these vulnerable association areas, may increase the susceptibility of cognitive cerebellar regions to age-related network disruption, whereas motor cerebellar regions, supported by more specialized network connections, appear better able to preserve functional resilience. Importantly, this account is tentative and requires further empirical validation to elucidate the mechanisms driving differential resilience of cerebellar function across functional domains.

### 4.4. Preserved cerebellar motor performance in aging due to cerebellar reserve or due to compensatory strategies?

An important question raised by the present findings is whether the observed resilience in cerebellar motor function reflects preservation of cerebellar-specific mechanisms themselves or whether it is supported by compensatory strategies.

One possible explanation for the absence of cerebellar motor function–structure associations is that the task outcome measures may not have been sufficiently specific to isolate cerebellar processing, allowing motor performance in older age to be supported by compensatory recruitment of additional cortical regions involved in motor and executive functions. However, in the current context, compensation via non-cerebellar networks appears less likely to support task performance efficiently, as the outcome measures were specifically chosen to isolate cerebellar-dependent motor mechanisms. Crucially, evidence from studies in patients with focal cerebellar damage performing comparable motor and timing tasks demonstrates marked impairments in these same performance parameters, despite largely preserved cortical motor function (Bastian et al., 1996, 2000; Franz et al., 1996; Ivry et al., 2002; Ivry & Keele, 1989). This indicates that intact cerebellar processing is necessary for successful task performance and that recruitment of other brain regions would therefore be insufficient to fully substitute for cerebellar function in these paradigms. Another possible compensatory strategy is that older adults recruit neural resources more strongly or earlier to maintain performance at lower task demands, a pattern described by the compensation-related utilization of neural circuits hypothesis (CRUNCH) (Reuter-lorenz & Cappell, 2008). Consistent with the CRUNCH framework, it remains possible that older adults maintain performance through increased recruitment within cerebellar motor regions, rather than through engagement of alternative cortical networks. In line with this interpretation, Bernard et al. (2020) reported that older adults more consistently recruited the same motor cerebellar regions across different motor tasks compared with young adults. This stable and repeated pattern of activation can be interpreted as a form of successful cerebellar scaffolding that preserves motor performance (Bernard et al., 2020).

Critically, if successful cerebellar scaffolding supports the maintenance of motor performance, this compensatory strategy may come at a cost: neural activation tends to occur earlier and in a less differentiated manner, potentially reducing the flexibility of available neural resources (Bernard et al., 2020; Reuter-lorenz & Cappell, 2008). Therefore, if task demands increase, the capacity limits of these neural resources may be reached earlier, making this form of compensation ineffective under higher task demands and leading to performance declines. This interpretation raises the possibility that the cerebellar-specific motor tasks employed in our study were not sufficiently complex to reveal age-related functional decline. The pure elbow motion and motor timing tasks involved relatively isolated upper-limb movements and enabled precise extraction of cerebellar-specific outcome measures. Although we aimed to challenge the motor system optimally by introducing standardized task-specific stressors, the inherent demands of both tasks on the motor system remained relatively low compared with more complex whole-body tasks. Consequently, cerebellar scaffolding may have effectively masked subtle age-related differences in the cerebellar processes that are critical for determining task performance.

In contrast, when considering more challenging whole-body movements, it becomes increasingly difficult to isolate the specific cerebellar contributions to task performance. In the present study, we investigated a postural balance task that imposed substantially higher demands on the sensorimotor system by simultaneously engaging multiple effectors and neural systems. Notably, clear age-related declines in balance performance were observed in this whole-body task. Although we classified the balance outcome measure as a general sensorimotor measure, balance control is known to emerge from dynamic interactions among multiple systems, including sensory, cerebellar, subcortical, cortical, and spinal mechanisms (Dijkstra et al., 2020; Surgent et al., 2019). Consequently, the observed age-related decline in balance performance is unlikely to reflect sensorimotor changes alone, but may also be influenced by age-related alterations in cognitive control (Bahureska et al., 2017) and/or specific cerebellar motor circuits. Under these challenging task conditions, it is therefore possible that cerebellar scaffolding can no longer fully compensate for age-related cerebellar degeneration, resulting in reduced adaptability and automaticity of balance responses in older adults. However, based on the present outcome measures, we are unable to disentangle the relative contributions of these neural mechanisms.

While it remains possible that some age-related effects were masked by limited task complexity, it should be noted that, although the pure elbow motion task involved only the upper limbs, the required movement speed was highly demanding and approached the maximum level feasible within an experimental setting. Even if this task did not fully challenge the older adults, it is noteworthy that the older-old adults, who would be expected to show the greatest age-related decline, were nevertheless able to maintain cerebellar-specific motor performance at a level comparable to that of young adults. Taken together, although task complexity may have attenuated the detection of subtle age-related differences in cerebellar motor outcomes, the preserved performance observed in the pure elbow motion task, particularly in older-old adults, supports the notion that cerebellar motor function can be maintained during healthy aging, potentially reflecting the presence of a cerebellar motor reserve rather than reliance on compensatory strategies alone.

### 4.5. Functional resilience emerges from context-dependent system stressor responses that cannot be generalized across the individual

One of the most striking findings of our study was the absence of correlations between resilience measures across all tasks, indicating that individuals’ resilience does not generalize across tasks or functional domains. Even between two cerebellar-specific motor tasks that showed relatively preserved performance in older adults, resilience measures were not significantly associated. Across all task pairings, correlation coefficients were uniformly low (all r < 0.30) and not significant, strongly suggesting that functional resilience is highly task-and domain-specific rather than a stable individual trait.

This absence of association challenges some aspects of complex systems theory, which conceptualizes humans as complex adaptive networks composed of interacting subsystems whose resilience is assumed to be coupled to that of the system as a whole (Scheffer et al., 2018). The lack of cross-task associations complicates the inference of a general, individual-level resilience and suggests limited coupling between functional subsystems. Notably, although such coupling is thought to become more apparent during aging, when increases in mortality risk are thought to reflect declining overall resilience (Scheffer et al., 2018), we observed similarly weak associations between resilience measures in older and younger adults.

At the same time, complex systems theory also acknowledges that age-related loss of resilience across the body is often heterogeneous, with different subsystems showing distinct trajectories. This heterogeneity may help explain why resilience responses to increased task demands were more pronounced for outcomes relying primarily on cerebellar motor mechanisms than for more general sensorimotor or cognitive outcomes. Although the possible associations between resilience responses across different neural mechanisms need to be better understood, our findings underscore the need for caution when inferring an individual’s overall resilience from that of a specific subsystem, as preserved function in one domain does not necessarily imply resilience to stressors affecting other domains or subsystems.

### 4.6. Limitations and future directions

This study has several limitations. First, the study design was not optimal for capturing within-person resilience. The tasks and stressors employed primarily quantified resilience in terms of the system’s immediate response to a stressor. Although standardized stressors are valuable for probing how a system reacts to challenges, other important dimensions of resilience, such as recovery dynamics and fatigability, could not be assessed within the applied task paradigms. In addition, the cross-sectional design precluded inferences about within-person trajectories of resilience over time. Longitudinal studies will therefore be required to determine how different dimensions of resilience evolve across the adult lifespan and whether individuals exhibit stable or changing resilience profiles during healthy aging.

Second, our MRI approach was limited to cerebellar grey matter volume and therefore did not capture the full complexity of brain function. Although grey matter volume is a widely used measure in aging research and provides information about the structural integrity of neural populations, it does not reflect the brain’s structural or functional connectivity. Future studies should therefore incorporate diffusion MRI and functional MRI to characterize connectivity within and between cerebellar functional regions, as well as their interactions with other task-relevant cortical areas. Such multimodal approaches would provide a more comprehensive understanding of the neural mechanisms underlying reserve and resilience in relation to the functional outcomes assessed here.

Finally, future research should focus on identifying potential protective factors that support resilience, as understanding which factors confer vulnerability or protection is essential for developing strategies to promote healthy aging. From a clinical perspective, the present findings suggest that cerebellar-mediated mechanisms may represent a promising target for interventions aimed at preserving motor function in older adults. At the same time, the task-and domain-specific patterns of resilience observed here highlight the importance of domain-specific assessment and rehabilitation strategies, rather than assuming generalized resilience across functional systems. Future work should therefore determine whether lifestyle or therapeutic interventions exert broader effects across multiple functional systems or remain largely confined to the specific neural circuits they directly engage.

## Conclusion

In conclusion, this study shows that cerebellar-dependent motor functions remain resilient across the adult lifespan, even under increased task demands and despite age-related structural decline within the cerebellum. This dissociation between preserved function and structural degeneration supports the concept of a cerebellar motor reserve that selectively contributes to the maintenance of motor performance in healthy aging. In contrast, general sensorimotor and cognitive functions exhibited reduced resilience, underscoring the heterogeneous nature of age-related changes across functional systems. At the same time, the absence of associations between resilience measures across tasks indicates that resilience is likely context- and function-specific rather than a stable person-level trait. Together, these findings advance a more nuanced understanding of reserve and resilience mechanisms in healthy aging. They emphasize the cerebellum’s distinctive role in sustaining motor function and underscore the need for integrative, multimodal research approaches to clarify how structural, functional, and behavioral factors interact.

## Acknowledgment

This work was supported by the Fonds Wetenschappelijk Onderzoek (FWO) (G095121N awarded to JJO).

